# Spatiotemporal patterns of cortical microstructural maturation in children and adolescents with diffusion MRI

**DOI:** 10.1101/2023.03.31.534636

**Authors:** Kirsten M. Lynch, Ryan P. Cabeen, Arthur W. Toga

## Abstract

Neocortical maturation is a dynamic process that proceeds in a hierarchical manner; however, the spatiotemporal organization of cortical microstructure with diffusion MRI has yet to be fully defined. This study characterized cortical microstructural maturation using diffusion MRI (fwe-DTI and NODDI multi-compartment modeling) in a cohort of 637 children and adolescents between 8 and 21 years of age. We found spatially heterogeneous developmental patterns broadly demarcated into functional domains where NODDI metrics increased and fwe-DTI metrics decreased with age. Using non-negative matrix factorization, we found cortical regions that correspond to lower-order sensory regions mature earlier than higher-order association regions. Our findings corroborate previous histological and neuroimaging studies that show spatially-varying patterns of cortical maturation that may reflect unique developmental processes of cytoarchitectonically-determined regional patterns of change.

## Introduction

Childhood and adolescence is a crucial period of extensive brain maturation, leading to major biological, cognitive and socioemotional transitions that have massive implications for health outcomes across the lifespan (1, 2). The neocortex constitutes the computational building blocks of the brain and is composed of structurally and functionally distinct cytoarchitectonic regions organized along broad functional gradients. Neocortical maturation is a protracted and asynchronous process that continues throughout childhood and adolescence and proceeds in a hierarchical manner, such that primary unimodal regions involved in sensorimotor processing develop earlier than higher-order association regions that subserve complex cognitive abilities (3). Therefore, understanding the microstructural trajectory of cortical maturation during this period can provide insight into not only the structural reorganization of the cortex during periods of heightened plasticity, but also the underlying mechanisms that contribute to cognitive and behavioral development.

Consistent with established histological timelines (4, 5), previous neuroimaging studies have found heterogeneous patterns of cortical maturation *in vivo* using MRI. Using morphological and volumetric analyses, earlier research efforts have uncovered asynchronous patterns of cortical thinning over the course of childhood and adolescence that is believed to reflect synaptic pruning (6, 7). More recently, others have characterized cortical microstructural maturation throughout development using more specific measures of intra-cortical myelination, including the ratio of T1w to T2w (T1w/T2w) contrasts (8, 9) and macromolecular proton fraction (10). In particular, Grydeland and colleagues assessed cortical T1w/T2w measures using growth modeling across the lifespan and found cortical myelination proceeds in successive waves that comprise a pre-pubertal wave of sensorimotor maturation and a post-pubertal wave of association and limbic maturation (9). While these studies provide insight into the important developmental process of cortical myelination, they lack sensitivity to the wide variety of cellular features that occupy the cortex, such as cell bodies, dendritic arborization and fiber orientations and there is a need to explore the developmental dynamics of cortical maturation using complementary measures of microstructure.

Diffusion MRI (dMRI) is an imaging technique that describes tissue properties by measuring water displacement patterns in the brain (11) Unlike semi-quantitative measures, such as T1w/T2w signal intensity differences, dMRI models yields biophysical quantitative values that correspond to histologically validated cellular features (12, 13). In particular, multi-compartment dMRI models provide enhanced specificity compared to more traditional diffusion imaging methods. This technique acquires diffusion-weighted images at multiple gradient strengths in order to probe tissue compartments with different diffusion profiles (14, 15). Neurite orientation dispersion and density imaging (NODDI) is one such model that characterizes restricted, hindered and free diffusion to model the intracellular, extracellular and CSF compartments separately to quantify neurite morphology (16). Additionally, free water-eliminated diffusion tensor imaging (fwe-DTI) (17) is an extension of the traditional DTI that models the tissue signal with an anisotropic tensor, but with an additional free water compartment to overcome partial volume effects with cerebrospinal fluid (CSF) along the pial border (18) and in perivascular spaces (19). Both NODDI and fwe-DTI provide complementary insight into brain microstructure as they model distinct aspects of molecular water diffusion. While DTI provide sensitive, but non-specific, insight into overall water diffusivity, NODDI is a biophysical model that quantifies parameters representative of known cellular phenomena (20).

dMRI is commonly used to study the maturation of white matter microstructure, due to the sensitivity of dMRI to the prominent orientational quality of myelinated fibers, and researchers have repeatedly shown heterochronous patterns of major white matter tract maturation that continue into the third decade of life (21–23). However, high-resolution diffusion-weighted images can also provide insight into cortical cytoarchitectonic properties (24). Previous studies have characterized the spatial gradients of high-resolution dMRI neurite metrics in the adult cortex, where they have found transitional boundaries in the dMRI cortical maps co-localize with areal boundaries of cytoarchitectonic regions derived from multi-modal segmentation and post-mortem methods (24–26). Additionally, known structural phenomena outlined in histological studies, such has hemispheric lateralization of cortical cell density, has been replicated in adults using NODDI (27). Age-related alterations to cortical dMRI metrics have been described in neonates and children as well. Significant decreases in DTI measures and increased NODDI measures have been described in neonates during the third trimester, and these developmental trends correlate with cortical thickness, cortical volume and associated white matter maturation (28, 29). Collectively, these rapid developmental alterations in the cortex *in utero* are believed to reflect increased cell density and dendritic arborization (28). Rapid alterations to cortical NODDI and DTI measures have also been reported postnatally across infancy and childhood (30). However, the developmental trajectory of cortical microstructure in children and adolescents has yet to be elucidated. It would be informative to explore how spatial patterns of cortical microstructure evolve in concert with periods of enhanced plasticity and rapid behavioral changes during this dynamic period of life, and possibly to help detect aberrations associated neurodevelopmental disorders.

The goal of the present study is to characterize the spatiotemporal development of cortical microstructural maturation in a large cohort of 621 typically developing children and adolescents between 8 and 21 years of age from high quality diffusion MRI dataset collected from the Human Connectome Project Development study. Cortical microstructural metrics derived from fwe-DTI and NODDI were sampled along mid-cortical surface vertices to assess local developmental alterations. We determined the main effect of age on regional cortical microstructure using a voxel-wise approach. We also characterized the magnitude and timing of age-related changes in microstructural parameters within distinct cortical regions using growth models and then clustered cortical vertices using non-negative matrix factorization (NMF) to understand how microstructure across regions co-vary with age.

## Materials and Methods

### Subjects

Neuroimaging data used in the preparation of this study were obtained from a cross-sectional sample of 621 (N F) healthy children and adolescents between the ages of 8 and 21 years (*M*±*SD*=14.36±3.77 years) recruited through the Lifespan Human Connectome Project in Development (31). The Lifespan Human Connectome Project in Development is an extension of the Human Connectome Project with the goal to characterize the structural and functional attributes of the normal developing brain. Subjects were recruited and scanned in Boston, Los Angeles, Minneapolis and St. Louis. Participants were excluded if they were born premature, require special educational services, had MRI contraindications, or had a history of serious medical problems, head injury, endocrine disorders, psychiatric disorders, or neurodevelopmental disorders. Because there is little consensus surrounding the term “typical development,” participant inclusionary criteria encompassed a wide variety of traits and behaviors, and subjects were excluded if they could not complete the study or was diagnosed with a disorder that may alter the course of brain development.

### MRI acquisition

Subjects were scanned on a Siemens 3T Prisma with an 80 mT gradient coil and a Siemens 32-channel Prisma head coil (32). T1w multi-echo MP-RAGE scans were acquired with the following acquisition parameters: voxel size=.8 mm isotropic, 4 echoes per line of k-space, FOV = 256 × 240 × 166 mm, matrix = 320 × 300 × 208 slices, 7.7% slice oversampling, GRAPPA = 2, pixel bandwidth = 744 Hz/pixel, TR = 2500 ms, TI = 1000 ms, TE = 1.8/3.6/5.4/7.2 ms, FA = 8 degrees. Motion-induced re-acquisition were allowed for up to 30 TRs. Multi-shell diffusion MRI (dMRI) scans were acquired with the following acquisition parameters: 1.5 mm isotropic voxel, TR=3.23 s, MB factor=4. dMRI data was collected using 185 diffusion-encoding directions split into two runs (92-93 directions per scan) and acquired twice with opposite phase-encoding directions (AP and PA), resulting in a total of 4 runs. Two shells (b=1500 s/mm^2^, 3000 s/mm^2^) were interleaved within each dMRI run with 28 b=0 s/mm^2^ volumes equally interspersed across all the scans.

### MRI data processing

dMRI data was processed using the HCP dMRI processing pipeline (Sotiropoulos et al., 2013). Motion-induced, eddy-current and susceptibility-induced artifacts were corrected using FSL EDDY (Andersson & Sotiropoulos, 2016). This method harnesses the complementary information provided by pairs of b=0 volumes with reversed phase-encoding directions to calculate the susceptibility-induced off-resonance field, which is then fed into a Gaussian Process predictor to correct for these distortions.

T1w data was processed with the HCP minimal preprocessing pipeline (Glasser et al 2013). Triangulated meshes of the pial and white matter cortical surfaces were extracted using Freesurfer 6 (http://surfer.nmr.mgh.harvard.edu/) and the cortex was parcellated into 74 bilateral regions according to gyral and sulcal boundaries defined by the Destrieux atlas (35). To project microstructural measures onto the cortical surface, the T1w and average dMRI b=0 images were aligned in native space using a rigid body transformation implemented with the ANTs software (36) and dMRI microstructural features were spatially normalized to the high resolution T1w space. We created population-averaged templates of the DTI and NODDI parameter maps and T1w images from our dataset using a high-dimensional spatial normalization workflow implemented in DTI-TK (37) and these templates were aligned to the IIT template (38). Then, MRI data from study participants were spatially normalized to the template to derive a population-average cortical brain surface that allows for one-to-one correspondence in the surface-based mapping analysis.

### Diffusion MRI models

The dMRI data was first denoised using a non-local means filter and microstructural dMRI parameters for two multi-compartment diffusion models were calculated using the Quantitative Imaging Toolbox (QIT) (39). The diffusion tensor imaging (DTI) parameters (40) axial diffusivity (AD), radial diffusivity (RD), mean diffusivity (MD) and fractional anisotropy (FA) were derived from free water-eliminated diffusion tensor imaging (fwe-DTI) (17) using iterative least squares optimization (18) – fwe-DTI is a bitensor model that separately estimates the free water compartment using an isotropic tensor and the tissue compartment using an anisotropic tensor. This approach can reduce partial volume effects due to CSF contamination, which is particularly important in the gray matter tissue adjacent to the pial surface. The NODDI parameters neurite density index (NDI), isotropic volume fraction (FISO) and orientation dispersion index (ODI) (16) were calculated using the spherical mean technique implemented with the QIT (41). We utilized models that separately model contributions of the free water and tissue water to reduce partial volume effects, which is particularly important in tissue boundaries, such as the pial surface.

### Surface-based mapping procedure

Our surface-based mapping procedure is similar to that used by (26) with an additional statistical refinement procedure to reduce the influence of outlier values due to mis-aligned tissue boundaries. For each subject, we computed to cortical microstructural parameters on the mesh vertices using a two-step procedure outlined in (42). First, at each vertex, we sample the diffusion parameters at 15 equally spaced points between the inner and outer cortical surfaces and calculated a weighted average using a Gaussian function centered at the midpoint of the path and scaled by the cortical thickness, i.e. where the weighting at the midpoint was 1 and the inner and outer cortical boundaries had a weighting of .05. By using a weighted average, this scheme is designed to isolate gray matter voxels by reducing the influence of voxels that are closer to tissue boundaries. Then, we employed robust averaging using the M estimator approach to exclude outlier values with a z-score threshold of 3 standard deviations from the weighted average calculation. In the resulting cortical microstructural maps, Laplacian smoothing was carried out with 30 iterations of lambda=1.2 subsequent to parameter mapping to accommodate possible misalignment of the cortical registration algorithm. Microstructural features were also averaged within the pre-defined cortical regions and visualized on the population-average cortical surface.

### Statistical analysis

The main effect of age on diffusion parameters after controlling for the effects of scanner site and sex was carried out at each vertex using a general linear model to test the linear and quadratic age effects with random field theory (RFT) cluster-wise thresholding using SurfStat implemented in Matlab (www.math.mcgill.ca/keith/surfstat). A height threshold of *p*<.001 and a spatial extent threshold of *p*<.05 were used to assess significance. Across all vertices, Bayesian information criteria (BIC) was used to determine the best fit model among linear, quadratic and growth models.

To quantify the magnitude and timing of cortical microstructural maturation, we fit the three-parameter Brody growth model using nonlinear least squares regression implemented in R version 4.1.3 to each of the 76 bilateral cortical regions of the form: Metric = b+a*exp(-k*age). We then defined peak maturational age as the age at 90% of the asymptotic value, as described previously in (21). Nonparametric bootstrap resampling with replacement (N=10,000) was implemented to obtain standard error and confidence intervals for the growth model coefficients and peak age estimates.

In an effort to identify patterns of age-related microstructural covariation, we used non-negative matrix factorization (NMF) to cluster the cortical vertices using the dMRI metrics. NMF is an unsupervised, multi-variate statistical learning technique that has been used previously to analyze structural neuroimaging data (Sotiras et al., 2015). The goal of NMF Is dimensionality reduction and feature extraction through decomposition of a data matrix into two smaller matrices with *k* factors, such that the linear combination of these smaller matrices reproduces the original matrix. In accordance with previous studies (29), we performed NMF with *k*=4 using a matrix constructed from all the fwe-DTI and NODDI parameters on the cortical sheet vertices as features in the model. We then averaged the dMRI metrics in each cluster and fit Brody growth curves and calculated the age at terminal maturation to identify the developmental timing of each group of regions.

## Results

Here, we explored age-related differences in quantitative markers of cortical microstructure using multi-parametric diffusion mapping that provides insight into diverse tissue properties. Furthermore, we used growth models to capture the developmental timing of regional cortical maturation from childhood through late adolescence. To develop a more precise understanding of inter-regional cortical developmental gradients, we employed an unsupervised clustering approach to identify regions that co-vary with age.

### Mean cortical parameter maps

Cortical microstructure was non-uniformly distributed throughout the cortex with varying spatial patterns among DTI and NODDI metrics (**Figure 1**). Cortical NDI was highest in the primary somatosensory and motor regions around the central sulcus, primary auditory regions around the posterior segment of the lateral sulcus and early visual regions in the medial occipital lobe, posterior cingulate, parahippocampal cortex and medial orbitofrontal cortex. Elsewhere, NDI values were generally low in other cortical regions, including many higher order association cortices. Cortical FISO had similar spatial patterns as NDI, though the regions with elevated FISO were far more restricted compared to NDI.

**Figure 1.**
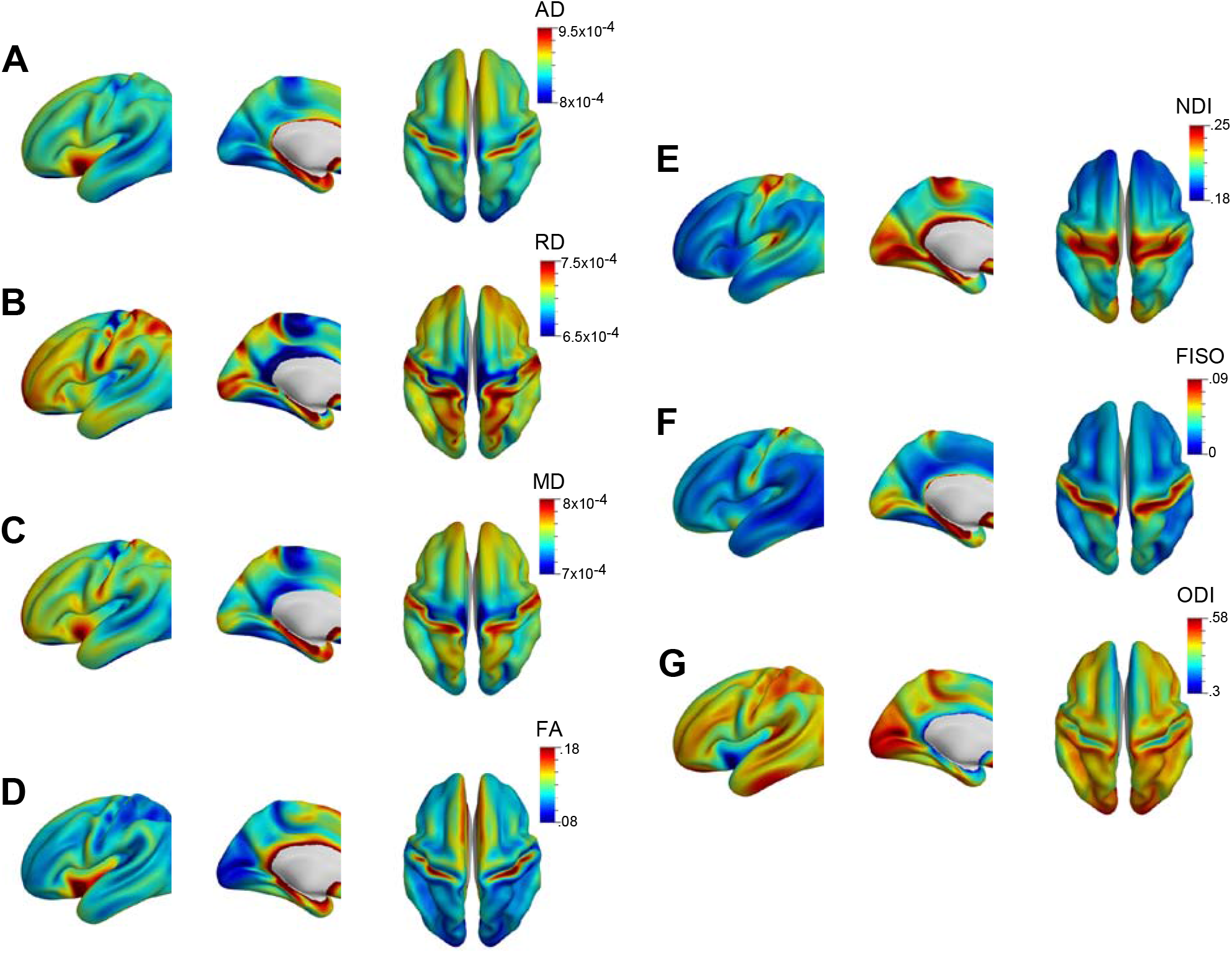
Group-averaged cortical surface maps for (A-D) DTI and (E-F) NODDI metrics.

Cortical ODI generally showed an inverse trend compared to NDI and FISO, where ODI was high throughout the cortex; however, similar to NDI and FISO, ODI was highest in the primary visual regions, primary auditory regions and medial orbitofrontal cortex. Unlike NDI and FISO, ODI was high in inferior temporal regions and was moderately elevated in precentral and postcentral gyri, but not the central sulcus. The distribution of cortical ODI generally aligned with the contours of the cortex, with elevated ODI in the gyri and lowest ODI in the sulci.

In general, DTI metrics showed similar spatial distributions throughout the cortex. Across all parameters, DTI metrics were highest in the parahippocampal region and early somatosensory regions adjacent to the central sulcus, and lowest in the early auditory regions around the lateral sulcus, medial occipito-temporal sulcus and paracentral lobule. Additionally, cortical FA, MD and AD was high in the insula. Some striking differences were also observed among DTI metrics. Cortical FA and AD was low in early visual areas of the cuneus and calcarine sulcus, while RD was high in corresponding regions. Cortical RD and MD was low, while cortical FA was high, in the posterior cingulate. Additionally, the superior parietal lobule had high RD and low FA, while the precentral gyrus had low RD and MD. Across all measures, the highest standard deviation in diffusion metric distributions was observed in the primary somatosensory regions near the central sulcus and medial orbitofrontal cortex (**Supplementary Figure 1**).

### Influence of age on diffusion metrics

Across the entire cortex, age was significantly associated with reduced mean AD (*r*(619)=-.63, *p*<.001), RD (*r*(619)=-.49, *p*<.001), MD (*r*(619)=-.55, *p*<.001) and FA (*r*(619)=-.45, *p*<.001). In order to explore the regional development of cortical tissue microstructure, we identified the main effect of age on DTI and NODDI measures, after controlling for the effects of sex and scanner type, using a vertex-wise approach with random field correction on the cortical sheet (**Figure 2**). Age was linearly correlated with reduced DTI metrics throughout the majority of cortical vertices, except for the superior parietal lobule and early visual areas around the cuneus and calcarine fissure. Additionally, age was not significantly associated with RD and MD within vertices of the somatosensory cortex, nor FA within primary motor and superior frontal regions. When testing for the quadratic effect of age on cortical microstructure, we found widespread nonlinear effects of age on cortical AD, RD and MD. Age-related alterations to cortical FA were best described with a linear model.

**Figure 2.**
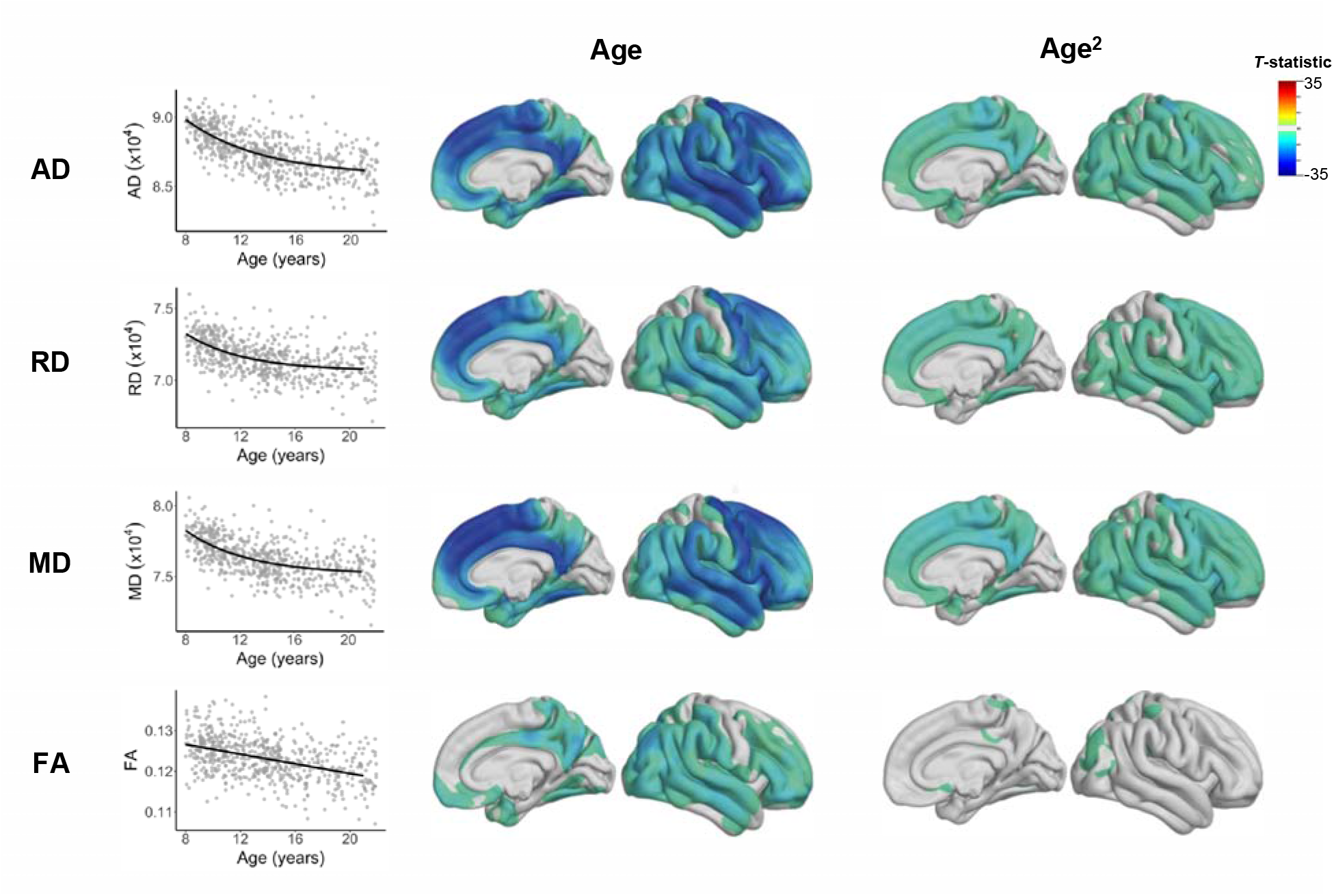
Main effect of age on cortical DTI parameters. (Left) The relationship between age and diffusion metrics averaged across all cortical vertices bilaterally is shown for AD, RD, MD and FA. Surface renderings represent t-statistics for the linear (center) and nonlinear (right) effects of age on DTI metrics after controlling for sex and scanner type. T-values are thresholded according to the significance level derived from random field theory. Overall DTI metrics decreased with age in the cortex. Using a vertex-wise analysis with significance threshold derived from random field theory, vertices that show significant linear (age) and non-linear (age2) effects of age on DTI metrics are shown after adjusting for sex and scanner type. Age is linearly associated with DTI metrics across the majority of the cortex, while non-linear age effects were broadly observed with cortical AD, RD and MD, but not FA.

Age was significantly associated with increased mean cortical FISO (*r*(619)=.48, *p*<.001), ODI (*r*(619)=.70, *p*<.001) and NDI (*r*(619)=.85, *p*<.001). Using the vertex-wise analysis after controlling for covariates, we found the influence of age on cortical microstructure varied across NODDI metrics (**Figure 3**). Age was linearly and nonlinearly associated with NDI throughout all cortical vertices. Cortical ODI was linearly associated with age throughout the majority of the cortex, with the exception of early visual regions that correspond to the calcarine sulcus and cuneus. Additionally, the lingual gyrus, isthmus of the cingulate gyrus, somatosensory cortex and medial orbitofrontal cortex showed significant nonlinear increases in ODI with age. While nonlinear associations between age and cortical FISO were not observed, FISO was linearly associated with age across large swathes of the cortex; however, age was not significantly associated with FISO in the temporal pole, superior temporal sulcus, anterior cingulate gyrus, superior frontal gyrus, medial orbitofrontal cortex and precentral gyrus.

**Figure 3.**
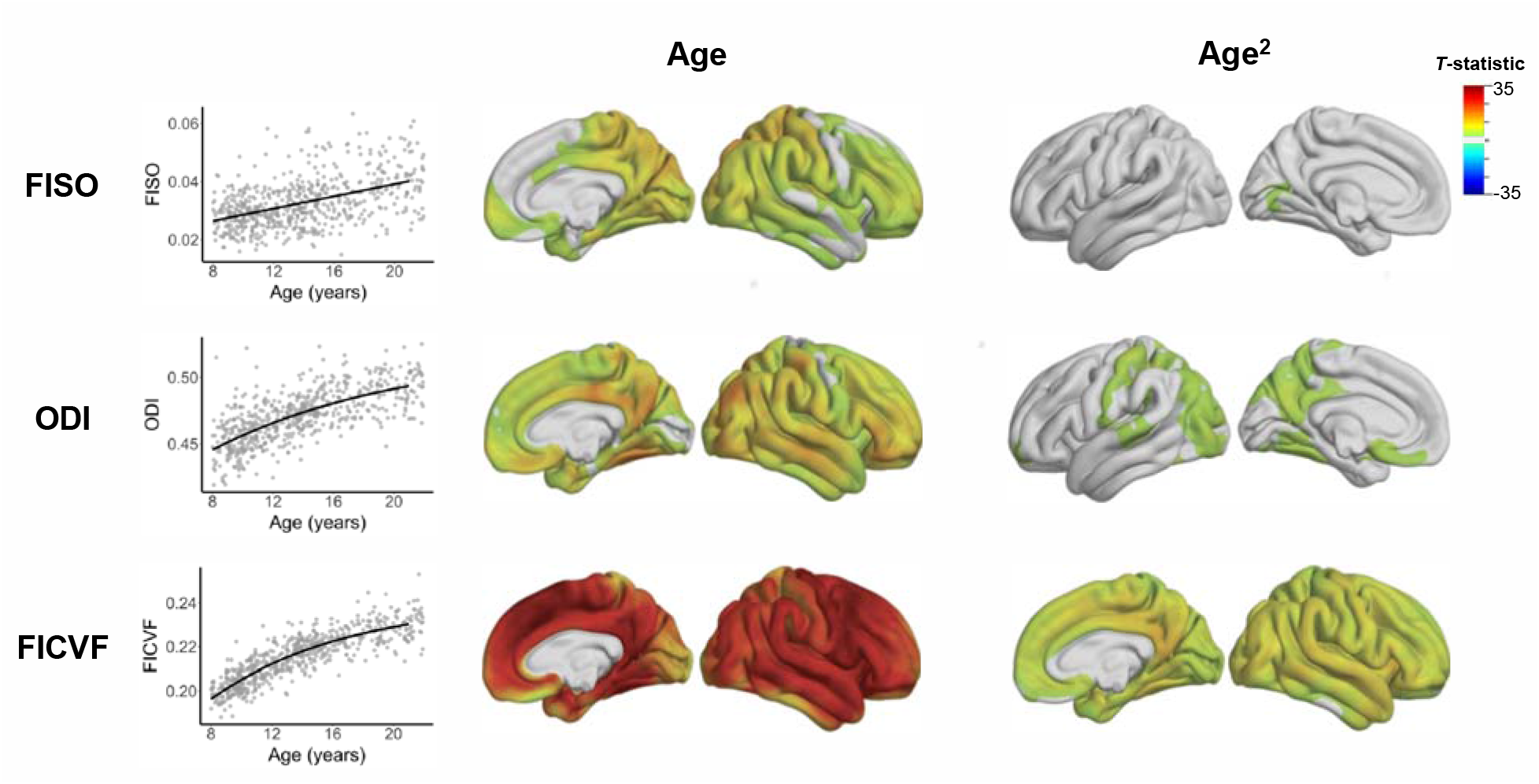
Main effect of age on cortical NODDI parameters. (Left) The relationship between age and diffusion metrics averaged across all cortical vertices bilaterally is shown for FISO, ODI and FICVF. Overall NODDI metrics increased with age in the cortex. Using a vertex-wise analysis with significance threshold derived from random field theory, vertices that show significant linear (age) and non-linear (age2) effects of age on NODDI metrics are shown after adjusting for sex and scanner type. Age is linearly associated with NODDI metrics across the majority of the cortex, whereas non-linear age effects were broadly observed with cortical FICVF.

### Time course of cortical microstructural maturation

In order to characterize the developmental timing of cortical microstructural maturation, nonlinear growth models were fit to diffusion parameters on the cortical surface and the peak age was defined as the age at 90% of the estimated asymptote. Overall, mean cortical NDI has a peak age of 15.63 [CI-95%: 10.37, 22.10]. The terminal maturational age for mean cortical AD, MD and RD is slightly younger, with estimated peak ages at 12.86 [9.59, 18.59] years, 10.58 [7.74, 15.56] years and 9.87 [6.90, 15.39] years, respectively. The growth curve produced unreliable estimates for the peak age of mean cortical ODI outside the sampled age range (25.58 [14.52, 61.89] years). Because nonlinear age effects were also not observed for FA and FISO cortical vertices, growth curves were not fit to FA, FISO and ODI in parcellated cortical regions.

The age at peak maturation for AD, RD, MD and NDI estimated within pre-defined cortical ROIs are shown in **Figure 4**. Across metrics, a general posterior-to-anterior pattern of maturation is observed, where occipital and parietal regions mature earlier than temporal and frontal regions. Individual cortical regions reach peak NDI between 9.62 and 25.57 years of age. AD, RD and MD mature across a broader age range, with individual ROIs reaching peak maturation between 2.89 and 30.41 years. Cortical microstructure for primary sensory regions tended to reach peak maturation earlier than other regions. The calcarine sulcus and cuneus, which includes the primary visual cortex, were among the earliest regions to mature for AD, RD, MD and NDI. Additionally, RD, MD and FA reached peak maturation earlier than surrounding regions within the primary auditory regions in the anterior transverse temporal gyrus. A striking demarcation is observed at the level of the central sulcus, where the postcentral gyrus reaches peak maturation earlier across microstructural metrics, including AD, RD and MD, compared to the precentral gyrus. Regions with protracted developmental timing across diffusion metrics include the anterior cingulate, superior frontal cortices, and the parahippocampal cortex.

**Figure 4:**
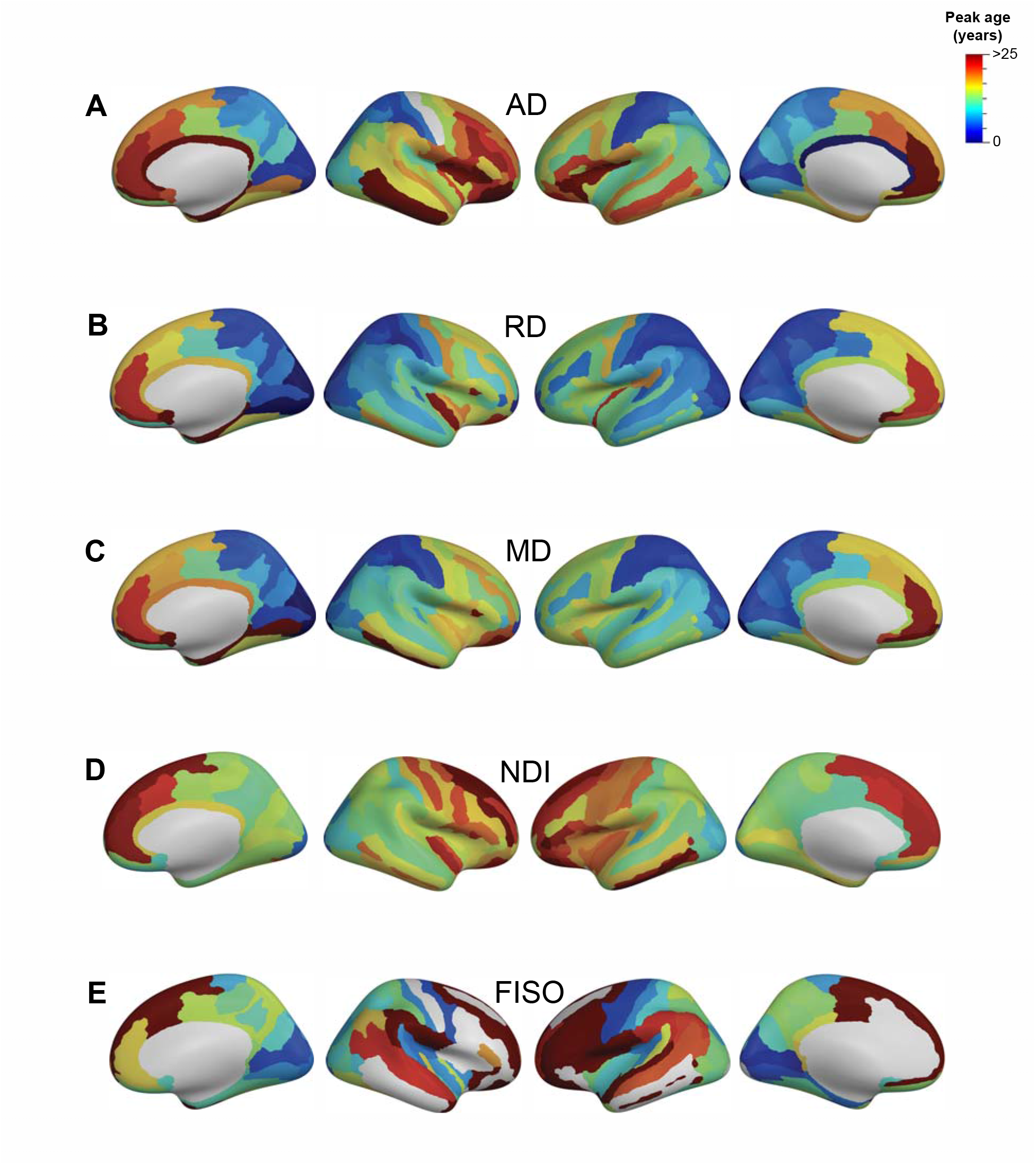
Developmental timing of cortical diffusion microstructure. The age at peak maturation, defined as the estimated age at 95% of the growth model asymptote, is shown within predefined cortical regions for diffusion metrics that underwent nonlinear age effects. The estimated age at peak maturation is shown for (A) AD, (B) RD, (D) MD, (E) NDI and (F) ODI. Regions where the growth model failed to converge and non-cortical surface points (e.g., subcortical midline) are shown in gray.

The correlation between age and microstructure as defined with FA, FISO and ODI also varies across the cortex (**Supplementary Figure 2**). FA and ODI showed the strongest correlations with age in regions that correspond to the middle temporal gyrus, posterior cingulate and precuneus. The strongest associations between FISO and age occurred in posterior regions, including the precuneus, superior parietal cortex, and postcentral gyrus. Interestingly, both ODI and FA showed the weakest correlation with age within the postcentral and precentral gyrus.

### Patterns of developmental covariation in cortical microstructure

To identify regions that demonstrate distinct patterns of developmental covariation, we used NMF to cluster cortical vertices according to their microstructural profiles across all subjects (**Figure 5**). This data-driven approach uncovered 4 clusters with common dMRI metric distributions: (1) A primary sensory cluster that includes the postcentral gyrus that corresponds to the somatosensory cortex, as well as the cuneus and pericalcarine cortices that correspond to primary visual cortex, (2) An occipito-parietal cluster that occupies the remainder of the occipital and parietal cortices, as well as the posterior superior temporal gyrus and precentral cortex, (3) A fronto-temporal cluster that consists of the frontal and temporal lobes, as well as portions of the supramarginal and inferior parietal cortex, and (4) a limbic cluster that consists of the parahippocampal cortex and the rostral anterior cingulate cortex.

**Figure 5:**
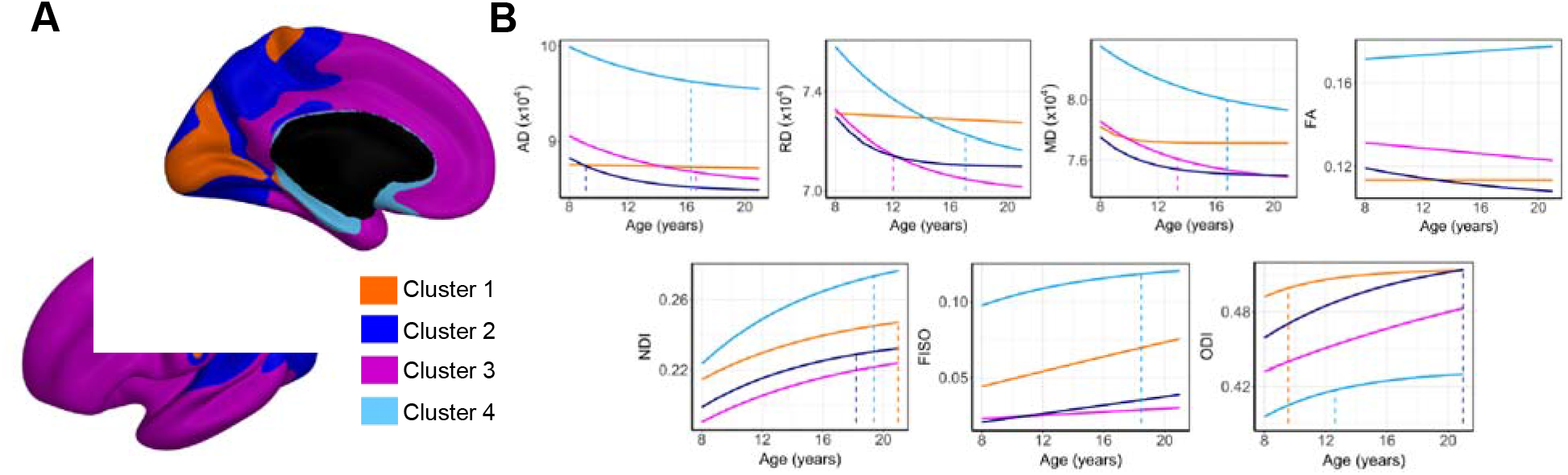
Cortical microstructural clusters derived from NMF. (A) Four clusters were identified using all diffusion parameters as features in the cortical mesh that correspond to primary sensory (cluster 1), occipito-parietal (cluster 2), fronto-temporal (cluster 3) and limbic cortices (cluster 4). (B) The relationship between age and each diffusion metric was assessed within each cluster and the line of best fit (solid line) and peak age estimate (dashed line) are shown for data best fit with a growth curve. Linear fits were found for cluster 1 (AD, RD, FA and FISO), cluster 2 (FA and FISO), cluster 3 (FA, FISO) and cluster 4 (FISO). Some terminal age estimates were observed outside the sampled age range and are not shown.

To better understand the developmental characteristics of the cortical clusters, the relationship between age and each diffusion metric was assessed with a Brody growth curve or linear model using Bayesian information criteria to determine the best fit line. Age-related alterations to FA and FISO within cortical clusters were best described with a linear model. FA was not significantly associated with age within the primary sensory cluster (p=.97). Age was associated with increased FA in the limbic cluster (*F*(1,619)=16.93, *p*<.001, *R*^*2*^=.03) and decreased FA in the frontotemporal (*F*(1,619)=166.5, *p*<.001, *R*^*2*^=.21) and occipitoparietal clusters (*F*(1,619)=-229.5, *p*<.001, *R*^*2*^=.27). Age was associated with increased FISO in the frontotemporal (*F*(1,619)=64.46, *p*<.001, *R*^*2*^=.09) and limbic clusters (*F*(1,619)=70.71, *p*<.001, *R*^*2*^=.10), and accounted for the most age-related variance in occipitoparietal (*F*(1,619)=223.9, *p*<.001, *R*^*2*^=.26) and primary sensory FISO (*F*(1,619)=225.6, *p*<.001, *R*^*2*^=.27). The developmental timing of NDI maturation was relatively consistent across cortical clusters, where NDI reached terminal maturation in late adolescence for the primary sensory (20.96 years), occipitoparietal (18.21 years), frontotemporal (23.95 years) and limbic (19.36 years) clusters. Across the remaining diffusion metrics, the primary sensory cluster reached peak maturation earliest. MD reached peak maturation by 3.23 years and ODI reached peak maturation by 9.55 years. Age was not significantly associated with AD (*F*(1,619)=1.76, *p*=19) or RD (*F*(1,619)=2.31, *p*=.13) in the primary sensory cluster. For AD, RD and MD, the occipitoparietal cluster reached terminal maturation by early childhood (AD: 9.12 years, RD: 6.06 years, MD: 6.92 years), followed by the frontotemporal cluster in mid-childhood (AD: 16.67 years, RD: 12.05 years, MD: 13.34 years), and then the limbic cluster by late adolescence (AD: 16.32 years, RD: 17.06 years, MD: 16.78 years). The pattern of maturation in ODI differed from the DTI metrics. ODI in the limbic cluster reached peak maturation by 12.61 years, followed by the occipitoparietal cluster at 20.96 years. Age was linearly associated with increased ODI in the frontotemporal cluster (*F*(1,619)=547.6, *p*<.001, *R*^*2*^=.47), where ODI was estimated to increase by .004 (SE=.0001) annually.

## Discussion

We have utilized developmental growth modeling and cortical region clustering to characterize the spatiotemporal pattern of cortical microstructural maturation from 8 to 21 years of age with multicompartment diffusion MRI models. We observed widespread and spatially-varying increases in fwe-DTI measures and decreases in NODDI measures throughout the cortex; however, different parameters yielded distinct developmental trajectories that may reflect unique neuroanatomical processes.

We observed a maturational gradient that proceeds in a posterior-to-anterior direction, such that occipital and parietal regions mature earlier than those in frontal and temporal regions across microstructural metrics. Our findings are in agreement with earlier studies on cortical gray matter development that have discerned a large-scale caudal to rostral direction of cortical maturation using measures derived from cortical morphology (6), T1w/T2w contrast (8, 9) and magnetization transfer imaging (10). Furthermore, this pattern of maturation extends beyond the cortex, as white matter exhibits similar developmental gradients where sensory pathways that project from occipital and parietal regions mature early in infancy (44), while maturation of frontotemporal tracts persist over the course of adolescence (45, 46). Architectonic cortical regions map to functional specialization in the cortex, and the posterior-to-anterior direction of cortical maturation corresponds to the spatial organization of the sensorimotor-association axis, as described by converging anatomical, functional and evolutionary data (3). Additionally, cortical maturation proceeds along the sensorimotor-association axis, such that lower order sensorimotor regions develop earlier than higher order association regions. Overall, our findings agree with this hierarchical model of maturation, as we observed occipitoparietal primary sensory regions mature by early childhood across diffusion metrics, while measures in frontotemporal transmodal regions reach adult-like levels by late adolescence and early adulthood.

Our findings of global increases in NODDI parameters over the course of development are in agreement with previous studies that found NDI and ODI in cortical parcels increased with age, though in a much younger sample of subjects between 0 and 14 years of age (30). The distribution of DTI and NODDI parameters on the cortex across participants was also largely in agreement with mean cortical parameter maps observed in adults (26). The spatial organization of NODDI metrics closely resembles the distribution of cytoarchitecture and myeloarchitecture from post-mortem brain samples (13), which suggests these diffusion metrics may reflect biological features. While the interpretation of diffusion contrast in white matter is relatively straightforward, diffusion contrasts in the cortex can be derived from several sources including axons, dendrites and cell bodies (47–49). Below, we attempt to disentangle the contributions of various cellular features to the diffusion metrics.

In the *ex vivo* rodent brain, cortical NDI was more strongly correlated with optical staining intensity of myelinated axons compared to non-myelinated axons (Jespersen et al 2010), which may be because myelinated axons restrict water molecule diffusion more strongly than non-myelinated axons (11). Additionally, the distribution of NDI is similar to histological myelin density in the spinal cord (20) and T1w/T2w contrast in the cortex (26). Together, these findings suggest cortical NDI is most sensitive to myelinated axons. However, since myelin content covaries with axon density (50) and neuron density (51–53), it is possible that variation in cortical NDI may be attributed to these other microscopic characteristics as well. ODI is sensitive to fiber heterogeneity, as shown in histology (20). Because axons preferentially align with radial or tangential axes in the cortex (12), variation in cortical fiber orientation dispersion may be influenced by the proportions of these fiber types. Additionally, dendritic arborization may contribute to orientational heterogeneity and neurite density in the cortex, though to a lesser extent than myelinated fibers.

Neuronal fibers also provide a source of anisotropic cortical diffusion contrast. Globally, we observed pronounced decreases in cortical AD, RD and MD and modest reductions in FA, which together suggests overall reduced rate of water diffusion due to the presence of molecular obstacles. In addition to molecular barriers introduced by neuronal axons, somata and dendrites (54), microglia and astrocytes may also obstruct water movement and influence the diffusion signal. Recently, researchers have demonstrated a correlation between microglial density and diffusion measures in in vivo and ex vivo animal models (55, 56). As systemic inflammation has been proposed to influence cognitive outcomes over the course of child development, this use of diffusion MRI as a proxy for cortical microglial infiltration may provide additional insight into health outcomes throughout childhood and adolescence. In another study carried out by Blumenfeld-Katzir and colleagues (2011), DTI was used to study the microstructural correlates of neuroplasticity in rats induced by spatial learning tasks. Authors found selective reductions in cortical MD in cortical tissue that were accompanied by enhanced immunohistochemical staining of synapses and astrocytes, but not myelin (57). While the effects observed in the study were transient, it is possible that the age-related MD reductions in the present study reflects long-lasting alterations in neuroplasticity due to astrocytic recruitment and synaptogenesis.

Regional variability in microstructural aging trends may provide insight into different cellular processes that govern cortical development in these regions. In particular, we observed a striking dissimilarity between the maturational trajectories of the primary sensory cortices and the primary motor cortex, particularly at the boundary between the postcentral gyrus (area 3b) and postcentral gyrus (area 4). Our clustering approach using NMF shows the postcentral gyrus and striate cortex develop earliest across all metrics. DTI measures were not significantly associated with age in these regions, which suggests that maturation of primary sensory regions is complete prior to 8 years of age. In contrast, the primary motor cortex showed significant age-related decreases in DTI metrics, but not NODDI metrics. The observed discrepancy in developmental patterns between sensory and motor regions may be attributed to the unique cytoarchitectural organization of these regions. Primary sensory cortex contain highly developed granule cell layers that consist of densely packed, small pyramidal and stellate neurons. Motor cortex, however, lacks clear granule layers and are defined by prominent large pyramidal cells. Age-related reductions to the overall ellipsoid value accompanied by increased intraneurite volume fraction in primary motor cortex may be due to increased pyramidal cell size over the course of childhood and adolescence. This conjecture is consistent with previous work that has found a protracted period of pyramidal soma enlargement in the cortex over the course of childhood and adolescence (58). The lack of DTI metric changes with age in primary sensory regions suggest the granule cell layer may be highly developed by late childhood, which is consistent with previous work that have found no developmental changes in granule layer morphology from young to adult mice cortex (59).

Developmental differences in the spatial distribution of cortical diffusion metrics may also be attributed to varying degrees of fiber organization and myelination in the cortex. Radial axonal processes from pyramidal cells constitute the dominant diffusion orientation in the cortex, particularly within the primary motor cortex (47, 60). Within the precentral gyrus, we observed significant age-related alterations in water diffusivity, but not orientation dispersion. Therefore, we hypothesize that maturation of the primary motor cortex during childhood is attributed to increased myelination of radial fibers in the cortex. Interestingly, we found the developmental trends in the precentral gyrus were in opposition to the postcentral gyrus, where orientation dispersion increased significantly with age, while water diffusivity remained constant over the course of development. The altered fiber orientation complexity in the postcentral gyrus may be due to increased myelination of tangential fibers with age. The inner and outer bands of Baillarger are dense tangential fiber bundles that traverse the internal granular layer and internal pyramidal layers, respectively, and is present in all cortical regions to varying degrees. The inner bands of Baillarger are enlarged in primary sensory cortical areas that receive extensive thalamic input and contributes to the prominent tangential diffusion observed in these regions (60). However, altered cortical ODI may also be due to technical limitations in our image acquisitions. Previous studies have shown that ODI is associated with reduced cortical thickness (26), potentially due to partial volume effects in the neighboring white matter and CSF compartments. Because the postcentral gyrus has overall thinner cortex compared to the precentral gyrus (61), it is possible that the increased ODI with age in the postcentral gyrus may be due, in part, to partial voluming in thin cortex with respect to the dMRI voxel size. However, we do not have sufficient resolution to distinguish the unique contributions of radial and tangential fiber types to the age-related diffusion signal alterations.

While the present study uses multiple diffusion metrics to provide complementary insight into diffusion properties across a large developmental age range, several limitations should be addressed. First, our voxel size cannot sufficiently distinguish cortical layers. Different layers are composed of distinct cytoarchitectonic populations that may differentially contribute to the diffusion signal. Furthermore, some diffusion metrics in thinner cortical regions may be obscured by partial volume effects, as described previously. Additionally, our youngest participants in the study are 8 years of age and we are therefore unable to track developmental timing in young childhood, where many cell populations undergo the most rapid growth. Lastly, the present study is cross-sectional and we are unable to make specific inferences about how diffusion properties change within children over time.

We used multiple diffusion metrics as surrogate markers for cortical microstructural maturation to quantify the spatiotemporal time course of the cortical sheet during typical development. Our findings support previous neuroimaging and histological work that demonstrates primary sensory regions mature earlier than higher order association regions. Additionally, we found differing developmental trajectories across NODDI and DTI metrics indicative of region-specific cytoarchitectonic patterns. The neurodevelopmental gradient maps developed in this study provides insight into neurotypical maturation and can be compared against patients with developmental disabilities to uncover alterations to the magnitude and timing of cortical maturation. The results from this study sheds new light on the maturation of cortical microstructure and demonstrates the utility of diffusion metrics to study the cytoarchitectural organization of the cortex. Future studies will aim to corroborate our findings with histologically matched templates to better understand how microstructural metrics derived from non-invasive neuroimaging techniques reflect the developmental processes that give rise to the emergence of cognitive functions and complex behaviors.

## Acknowledgements

The image computing resources provided by the Laboratory of Neuro Imaging Resource (LONIR) at USC are supported in part by National Institutes of Health (NIH) National Institute of Biomedical Imaging and Bioengineering (NIBIB) grant P41EB015922. Author KML is supported by the National Institute on Aging (NIA) of the NIH Institutional Training Grant T32AG058507. Author RPC is supported in part by grant number 2020-225670 from the Chan Zuckerberg Initiative DAF.

## Author contributions

<u>Kirsten M. Lynch</u>: Conceptualization, Methodology, Formal analysis, Investigation, Writing – Original Draft, Writing – Review and Editing, Visualization; <u>Ryan P. Cabeen</u>: Conceptualization, Methodology, Software, Data Curation, Writing – Review & Editing; <u>Arthur W. Toga</u>: Resources, Writing – Review & Editing, Supervision, Funding acquisition.

