## Supplementary Figure 2 for "Spatiotemporal patterns of cortical microstructural maturation in children and adolescents with diffusion MRI"

**Supplementary Figure 2**: Associations between age and diffusion metrics that undergo linear age-related alterations. The Pearson’s correlation coefficient, r, for the association between age and diffusion metric values within parcellated cortical regions is shown for FA, ODI and FISO.
